# Signatures of remote planning in hippocampal replay

**DOI:** 10.64898/2025.12.02.691753

**Authors:** Brian Lustig, Yingxue Wang, Sandro Romani, Eva Pastalkova, Albert K. Lee

## Abstract

During brief, intermittent “replay” events, hippocampal activity can express navigational trajectories disconnected from both when and where they originally occurred. While replay biased toward immediate future goals has been observed, there is no evidence yet linking replay to planning beyond the next action. Here, we designed a sequential spatial working memory task which required rats to utilize information across multiple temporally separated actions. Remote replay events matched the animal’s future navigational choices made after completing an intervening subtask. Critically, this occurred only when the replayed information was useful for reducing memory load, consistent with it being an active process. Our findings suggest these remote replay events are a neural correlate of episodic forethought, allowing animals to use memories to plan beyond their immediate surroundings.

## INTRODUCTION

One of the most powerful features of episodic memory is the ability to internally reproduce past experiences detached from both when and where they originally occurred (*1–6*). A major evolutionary advantage of being able to engage in such “mental time travel” may be its utility for memory-based planning (*4–7*) – for instance, thinking about where you could park your car if you were to visit your favorite restaurant later in the day. In this way, plans can extend beyond what is immediately in front of you and instead be made for situations that are spatially and temporally remote.

The hippocampus is known to be critical for episodic memory, spatial memory, and imagination (*2–4*, *7–9*). Previous work has found neurons in the rodent hippocampus that increase their activity during visits to specific locations in space (place cells) (*10*) or at specific moments in time (episode or time cells) (*11*, *12*); thus, any given experience is associated with a sequence of activated hippocampal neurons. Replay of these neural sequences was initially observed during population bursts associated with sharp wave ripples during sleep (*13*, *14*), which have been linked to memory consolidation (*15–17*). The subsequent discovery that replay also occurs in the awake state (*18–50*) opened up the possibility that it may function as a form of episodic memory retrieval and play a role in planning.

Awake replay has been linked to planning in studies where replayed trajectories are biased towards goals or future navigational choices (*24*, *29*, *32*, *36*, *38*) (but also see (*30*, *31, 34*)). However, those replays were not spatiotemporally remote, as they were associated with the animal’s next action and generally limited to regions visible to the animal. Importantly, awake replay can reflect spatially and temporally remote experiences (*21*), and thus represents an opportunity to capture evidence of internally generated, mental time travel-like activity in the service of planning, but such remote replay has not been previously linked to any behavior—including planning. To address this, we designed a working memory task that requires animals to make choices using spatially and temporally remote information and investigated the potential role of remote hippocampal replay in planning future behavior.

## RESULTS

We developed a self-paced memory task requiring visits to a repeating sequence of spatial goals to obtain reward (three-arm delayed sequence, or TADS, task). The experimental apparatus involved a maze consisting of three long arms with high walls that are connected to a central hub containing a running wheel (Fig. 1A). Rats needed to visit water ports at the end of each arm in the following order: Left (L) → Center (C) → Right (R) → Center (C) → repeat (Fig. 1B, top). Crucially, trials were divided into distinct spatiotemporal segments by a wheel run subtask between arms (*11*, *51*). At the start of each trial (Fig. 1B, bottom), rats traveled from the end of an arm to the central hub (segment 1, inbound), then the door closed, confining them there. To proceed, rats were required to complete a continuous eight-second wheel run in a given direction (which was the same direction for all trials, i.e. always facing to the left) above a predefined speed threshold (segment 2, wheel run). Upon successful completion of the wheel subtask, all three arm doors reopened, and the rat could select the next arm from among the three options (segment 3, outbound). A choice was registered by crossing a beam break near the end of an arm and, if correct, water reward was delivered (segment 4, reward consumption) (see supplementary movie S1).

**Figure 1.**
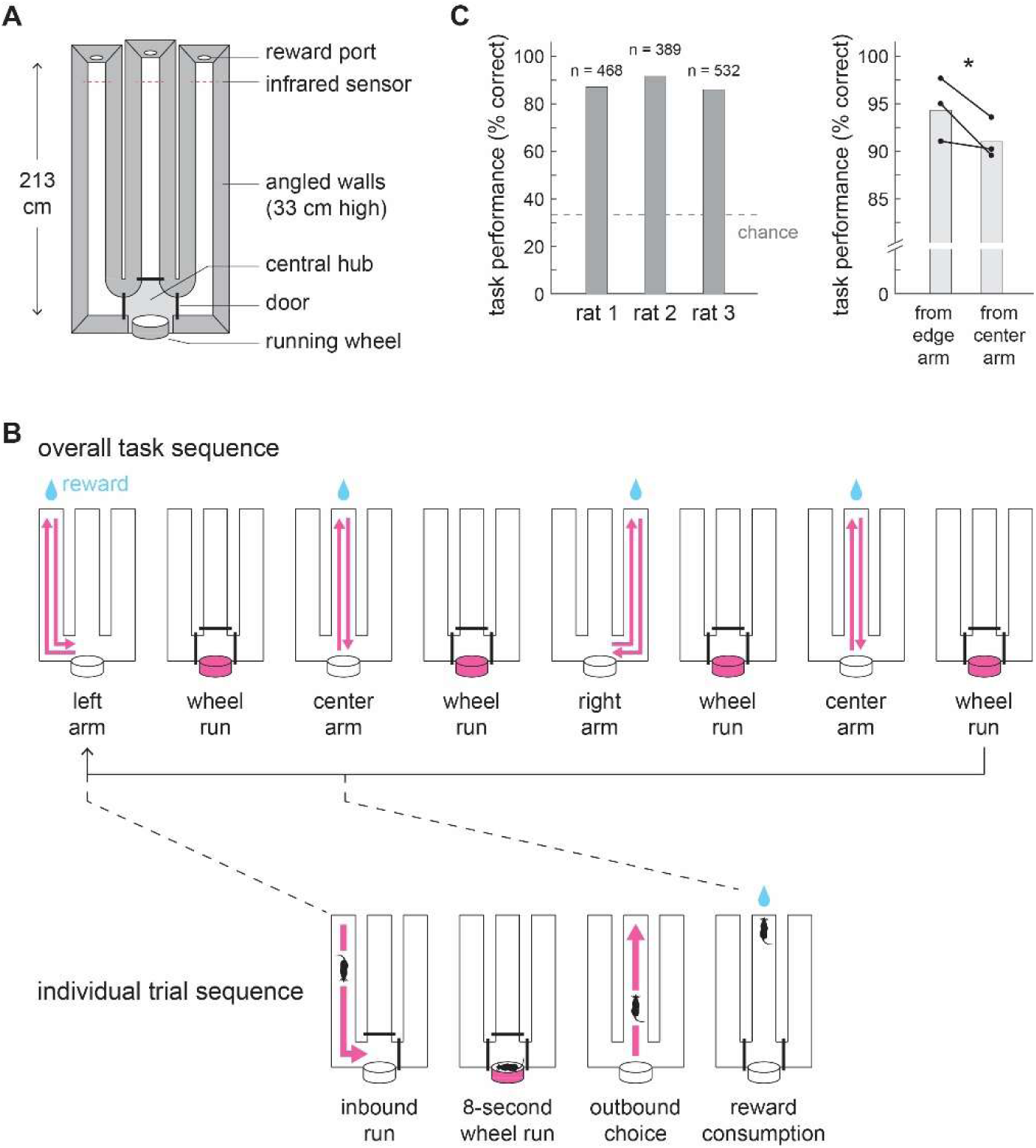
The three-arm delayed sequence (TADS) task. (A) Maze apparatus with high walls that visually isolate the three arms from each other. Central hub doors are automated. (B) Top: Illustration of correct sequence of arm choices, each separated by wheel run subtask, required to obtain reward at end of each arm. Bottom: Details of the multiple steps making up each trial. The inbound run starts at the end of the previous trial’s reward consumption. The wheel run subtask requires running continuously for 8 s above a threshold speed, after which all three central hub doors open, allowing the rat to make an outbound choice. If the correct arm is chosen, reward is delivered and the consumption period is the last phase of the trial. (C) Left: Mean task performance per animal (bars) (n = total number of trials per animal across sessions). Right: Mean task performance for trials beginning from an edge arm or the center arm (points and lines: per animal; bars: pooled across animals). Note that the edge arm performance considers only trials that start from a correctly chosen arm, and the center arm performance considers only trials in which the current and previous choices were correct, thus these values are higher than the overall mean task performance (see methods). *P < 0.05 (pooled).

All rats reached performance levels well above chance (mean: 86.1-91.9% of trials correct; N = 5, 4, and 6 sessions for rats 1-3, Fig 1C, left). Each full trial, from departing the current arm to the end of reward consumption, lasted 42.4 ± 18.3 s (mean ± s.d.), with durations for the behavioral segments of 5.1 ± 5.8 s (inbound run), 14.3 ± 10.9 s (hub/wheel run), 3.7 ± 3.9 s (outbound choice/run), and 19.2 ± 12.4 s (reward consumption). Importantly, there was a difference in cognitive load between trials starting from the center arm and those starting from either left or right arms (i.e. an edge arm). Specifically, for edge-to-center trials, the correct choice after the wheel run is the center arm, and this only requires remembering the inbound arm from before the wheel run (“1-back memory”). In contrast, center-to-edge trials require animals to also recall which outer arm it visited on the previous trial (“2-back memory”) and choose the opposite side. Consistent with differences in the cognitive demands of the two trial types, our task showed a small but significant difference in performance between them (edge-to-center trials: 94.3% correct, center-to-edge trials: 91.0%, *p* < 0.05, Fig. 1C, right).

We then examined hippocampal CA1 spiking activity recorded during the task to understand how neural dynamics support behavior. CA1 exhibited spatially and temporally selective firing across all segments of the task, such that sequences of directionally tuned place fields covered inbound and outbound arm run segments, and temporally tuned episode fields spanned the wheel-running period (Fig. S1). We applied established memoryless Bayesian population decoding methods (*20*, *24*) to the activity of all recorded cells to estimate the instantaneously represented location in the maze (and time during the wheel run) across all task segments. Figure 2A-D shows the behavior, corresponding neural activity, and decoder output for four consecutive trials. Following previous work on awake replay, we used the maps constructed from place and episode field firing to decode the brief ∼100 ms population burst events (PBEs; Fig. 2C,D) observed during reward consumption at each arm end. Decoding within PBEs was performed using a 20 ms sliding window stepped every 5 ms. PBEs showing a strong correlation between decoded locations and time—indicative of a smooth spatiotemporal progression—were classified as “trajectory replay events” (also referred to below as “replays”) (see Methods, Fig. 2D,E).

**Figure 2.**
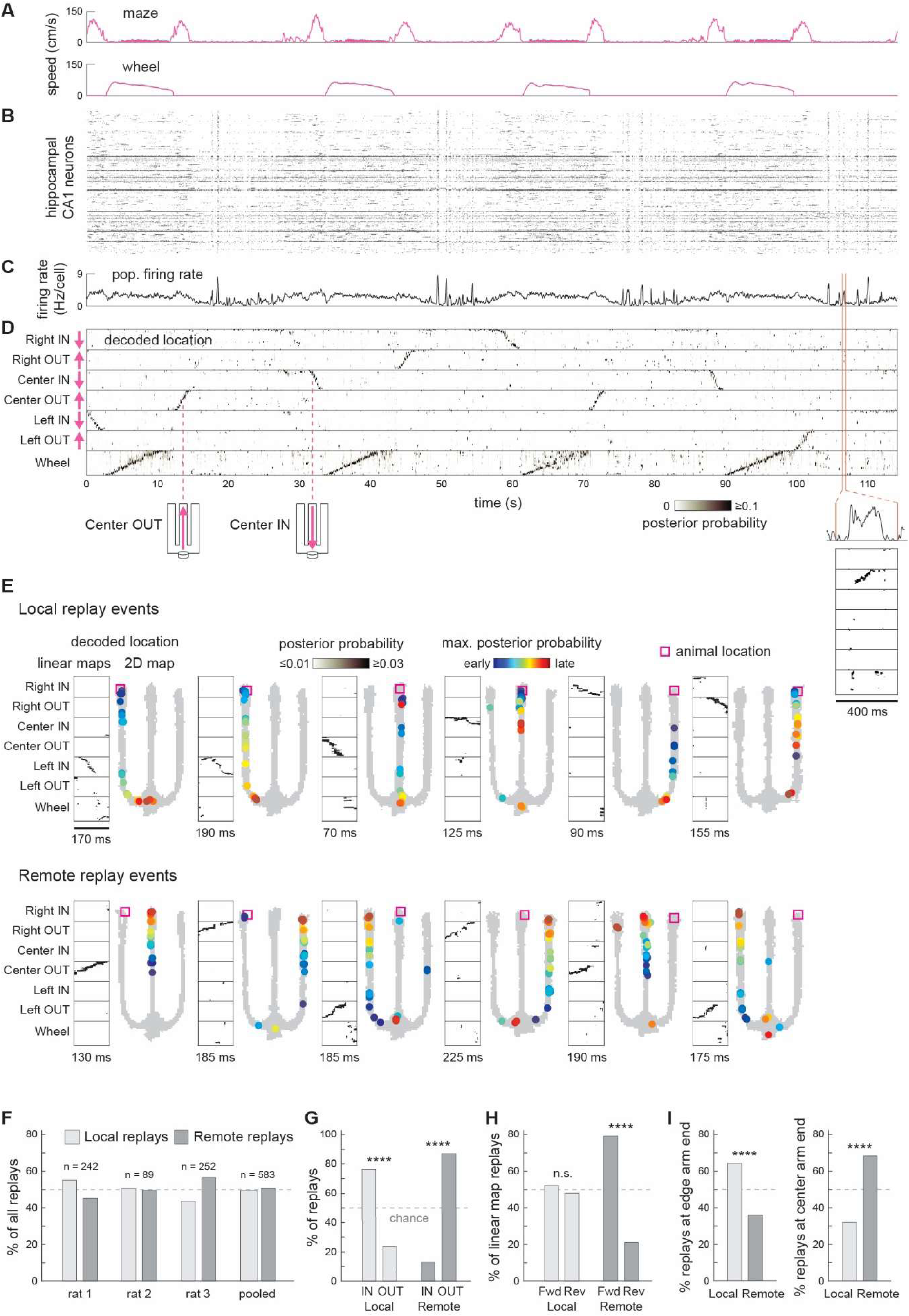
Hippocampal remote replays are strongly biased toward representing forward, outbound spatial trajectories and are more prevalent at the center arm end. (A-D) Example of four consecutive trials in the TADS task. (A) Animal movement speed in the maze and wheel. (B) Simultaneously recorded spiking activity of 315 dorsal hippocampal CA1 neurons. (C) Aggregate firing rate across all the neurons (1 ms bins, Gaussian smoothing with σ = 200 ms). (D) Memoryless Bayesian decoding of location onto seven linear maps derived from the tuned place field activity in inbound and outbound traversals of the three arms and the tuned episode field activity during the 8 s wheel run (20 ms bins, 5 ms step). Inset in (D) shows 400 ms of decoded activity during reward consumption at the left arm end, revealing remote replay of a spatial trajectory heading outbound in the right arm. This replay occurs during a population burst event (PBE) from (C) (inset: 20 ms bins, 1 ms step). (E) Examples of decoded PBEs (duration of each PBE shown below) that contained a qualifying replay event representing local (same arm as animal is in, top) or remote (different arm, bottom) replay trajectories (20 ms bins, 5 ms step) (see Methods). In addition to the seven linear maps as in (D), decoding was also done onto the 2-D map of the maze. (F) Proportion of all replay events that were local or remote (n = total number of replay events). (G) Proportion of local and remote replays with an absolute direction heading inbound or outbound. (H) Proportion of local and remote replays of one of the six linear maps that corresponded to a trajectory advancing in the forward running direction. (I) Proportion of local or remote replays occurring when the animal was consuming reward at an edge arm end or at the center arm end. (G-I) Results pooled across animals. ****P < 1 × 10^-6^, n.s. P > 0.05. Individual animal data consistent with pooled results displayed in Fig. S2.

We next characterized the content of these replay events. Awake hippocampal replay events have previously been reported to have a strong localization bias, whereby the trajectories tend to originate from locations where the animal currently is (e.g., (*18*, *19*, *24*)). While some studies have observed replay starting and ending away from the animal but spanning locations within the visible environment (*20*, *23*, *38*), few have investigated awake replay of locations that are spatially and visually isolated from the animal’s current surroundings (*21*, *36*). Here, we classified trajectory replay events into two categories—local and remote—based on the relationship between the replayed maze segment and the rat’s current location. Local replays were those that replayed locations in the currently occupied arm, which the rat had just traversed and would soon traverse again, reflecting proximity in both space and time (Fig. 2E, top). In contrast, remote replays depicted trajectories along one of the other two arms (Fig. 2E, bottom). These arms were neither visible from the rat’s current location nor immediately accessible, as reaching them required first completing the intervening wheel run task; therefore, remote replays reflected spatially and temporally distant, behaviorally disconnected representations. Both local and remote trajectory events were observed across all animals (Fig. 2E,F). Approximately half of all replays were remote (50.6% of n = 583 replay events across rats 1-3) (*21*). There was a clear difference in the preferred direction of propagation for local versus remote replay trajectories. Local replays showed a strong preference (76.4%, *p* < 1 × 10^-6^) for originating near the rat’s current location and propagating inbound toward the central hub, while the vast majority (87.1%, *p* < 1 × 10^-6^) of remote replays moved outbound toward another arm’s reward port (Fig. 2G). Most (79.0%, *p* < 1 × 10^-6^) of these remote outbound replays represented forward trajectories, i.e. they matched the sequence of place cells activated when the animal ran outbound to the reward on an arm (Fig. 2H). Interestingly, when the animal was in an edge arm, significantly more replays were local (64.0%, *p* < 1 × 10^-6^), while in the center arm, significantly more replays were remote (68.0%, *p* < 1 × 10^-6^) (Fig. 2I), suggesting the animal could be using remote replay to help perform the more demanding center-to-edge trials. (See Fig. S2 for individual animal and additional data.)

We then examined whether remote replays could reflect mental time travel-like planning for selecting the next arm, focusing on trajectory events occurring while the rat was located at the center arm reward port. This location provided advantages for assessing a possible functional role for remote replay. First, both remote arms (i.e. the left and right arms) had been rewarded in a balanced manner, reducing the potential influence of value or other bias on replay content (unlike for edge trials in which the center arm had been rewarded twice as much as the other edge arm) (*28*, *31*, *50*). Second, as described above, at the center arm end, animals needed to remember the previously chosen arm to correctly alternate to the other (unlike at the edge arm ends, see below). Finally, at the center location, the previously visited arm and the upcoming choice were distinct, allowing remote replay content to be linked to either the past or future (versus at the edge arms, where the center arm was both the correct previous and future choice). Figure 3A shows two examples of remote replays—one depicting the left arm, the other the right—while the rat remained stationary at the center reward port. In both examples, the remote replay matched the future arm choice. Importantly, these choices were made after the intervening wheel run subtask, ∼20s after the replay event (as illustrated in the time course of the behavioral trajectory). Strikingly, remote outbound replays (note again that the vast majority of remote replays were outbound, Fig. 2G and Fig. S2A) from the center showed a clear bias, significantly more often depicting the rat’s future behavior (82.7%, 84.2%, 74.3% for rats 1-3, mean 78.6%, *p* < 1 × 10^-6^, Fig. 3B,C; Fig. S3 shows that this result is robust to a range of replay detection parameter choices) (supplementary movie S2). Furthermore, these remote outbound replays generally represented trajectories in the forward direction (i.e. outbound replay in the outbound map for that arm), consistent with planning a run toward the next edge arm reward port (82.9%, *p* < 1 × 10^-6^, Fig. 3D) (*26*, *32*). These results strongly suggest that hippocampal replay can represent future-directed planning for spatially and temporally remote actions.

**Figure 3.**
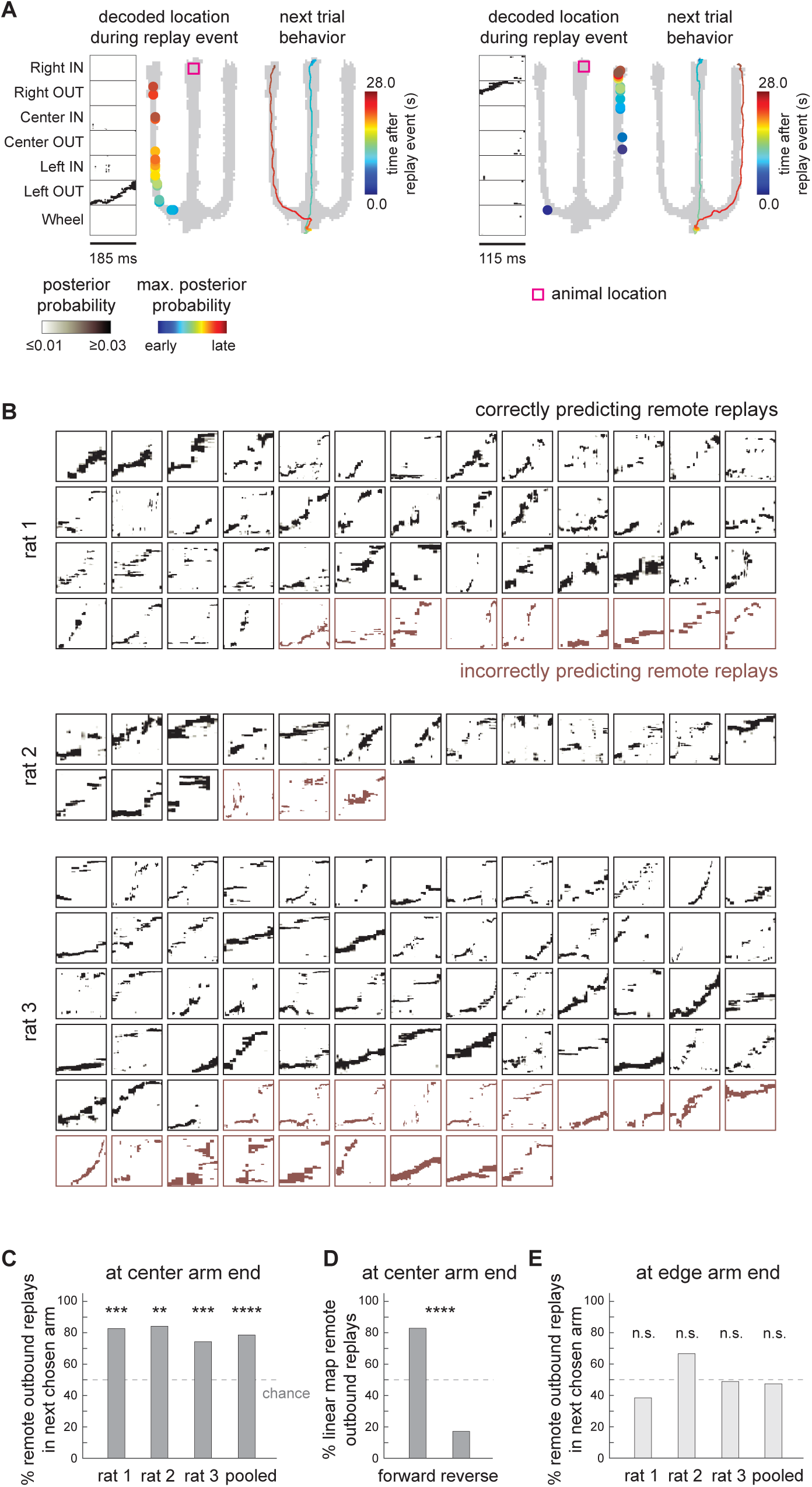
Remote replay at the center arm end is consistent with planning the future arm choice to be made after the intervening wheel run subtask. (A) Two example remote replays while the rat is at the center arm reward port that predict the subsequent behavioral choice. Note that, in both examples, rats only began the center arm inbound trajectory ∼10 s after the remote replay event, completed a >8 s wheel run, then chose the next arm ∼20 s after the event. (B) All remote outbound replays when the rat was at the center arm end. Replays correctly matching the rat’s next choice (black), and those not matching the next choice (red). Note that the vast majority of remote replays at the center arm end were outbound (Fig. 2G). (C) Proportion of remote outbound replays at center arm end that correctly matched the rat’s subsequent choice. (D) Proportion of center-arm remote outbound replays (of one of the six linear maps) that represented a trajectory advancing in the forward running direction. (E) Proportion of remote outbound replays from either edge (left or right) arm end that correctly matched the rat’s subsequent choice. Note that, unlike trials starting at the center arm, trials starting at an edge arm do not require the animal to maintain any information or plan their next choice while at that arm end (see text). **P < 0.01; ***, P < 0.001, ****P < 1 × 10^-6^, n.s. P > 0.05.

Lastly, we asked whether there was evidence that future-oriented remote replays did not just reflect passive mechanisms. Unlike periods at the center reward port, periods at edge arm ports do not require animals to maintain any memory to perform the upcoming edge-to-center trial. This is because the turn toward the central hub at the end of an inbound edge arm run provides a physical reminder that it is an edge-to-center trial (as do the distal visual cues above the maze walls), while the inbound center arm path in center-to-edge trials is by definition the same regardless of whether the previous trial was a left or right arm one. Therefore, if the observed bias for future-oriented remote replays at the center is not present in the edge arms, this would be consistent with center replays reflecting an active process of planning, as opposed to a passive mechanism that automatically replays the next arm in all conditions. Indeed, we found remote outbound replays at the edge reward ports showed no bias for either the future (center) arm or other arm (future arm proportion: 47.4%, *p* = 0.6, Fig. 3E).

## DISCUSSION

A major goal of neuroscience is to understand cognitive processes that go beyond stimulus-response relationships – for instance those that use internal models to enable complex behavior. Recalling memories of past experiences for the purpose of planning the future is an example of such a process. Here, we found hippocampal replay that matched the animal’s future path to a remote location, appeared well before execution (and before an intervening subtask), and occurred at a point (the center arm end) in a multi-step task where planning this action would help to reduce an otherwise high memory load. This strongly suggests that these remote replay events are neural correlates of planning and provide a specific example of episodic forethought (*4–6*).

The function of hippocampal replay has been a topic of much investigation, largely because of the high signal-to-noise ratio of the sequential activity patterns they display. Replay (and the PBEs during wake and sleep of which trajectory replay events are a subset) has been clearly implicated in learning and memory consolidation (*14–18, 22, 34*, *35*, *37, 40–50, 52, 53*), but investigation into the role of awake replay in planning has been characterized by varied results (*49*, *50*). Studies have shown that features of awake replay can predict whether future actions will be correct or not (*25*), and that prolonging replay event durations improves task performance (*42*), while disrupting them impairs it (*40–43*) (but see (*44*)). However, studies showing replay trajectories predicting future actions (*24*, *32*, *36*) have been contrasted by ones in which avoided (*30*), non-chosen (*31*), or past (*34*) routes are replayed. One possibility is that the function of replay depends on the specific design and demands of each task. In line with this, studies using the standard version of the three-arm alternation task, which has no running wheel or other subtask, have so far not reported a bias for future– or past-oriented replay while at the center arm end. The addition of an intervening subtask in the present study may be responsible for the difference because the wheel run makes the previous arm and future decision more distant in time, which can therefore be countered with replay-based planning of what to do next. Specifically, in center-to-edge trials, if the animal were to decide the next arm choice after the wheel run without planning, this would require using 2-back memory: memory of the arm it just came from (the center arm) as well as of the previous arm (one of the edge arms). However, by planning the next choice at the center arm end, the animal only needs 1-back memory (i.e. memory of the plan) after it exits the wheel. Note that making the plan itself at the center arm end also only requires 1-back memory (remembering the previous arm). In contrast, in the standard three-arm alternation task, the next arm choice in center-to-edge trials only ever requires 1-back memory.

Our findings build on previous work showing future-oriented replay in several ways. First, the spatiotemporal remoteness of task-relevant replay shown here has not been seen before. In other studies, the future path was associated with the next action and generally visible (*24*, *29*, *32*, *38*) (though not in (*36*)). Furthermore, in the latter case (*36*), the future-oriented replay was present during training but not in the trained condition, while the replay seen here is in a highly familiar task. Second, the future-oriented replay we observed at the center reward port took place under the conditions of a balanced, multiple-(here, two-) choice working memory task, while previous work involved a single goal (*24*, *36*) or involved multiple choices under a range of conditions (*32*). Third, as described above, we found future-oriented replay at the center but not edge arms, which serves as an internal control suggesting that the future replay at the center reward port is part of an active planning process rather than a passive process. Notably, recent work (*54*) has shown that the content of replay could at least partially be explained by a passive mechanism in which replay avoids the most recently activated locations (*54*, *55*). However, our future-predicting replay involves remote trajectories that are outside the proposed adaptation time window. Furthermore, remote replays at the edge arms were not biased away from the center (more recently experienced) arm.

The idea that animals can actively plan their future behavior by replaying specific trajectories agrees with recent hippocampal brain-machine interface (BMI) experiments (*56*, *57*). Those studies demonstrated that animals can volitionally control the activation of non-local hippocampal spatial representations, which is a fundamental process enabling mental time travel, imagination, mental simulation, and planning. However, a question that arises from this work is when do animals voluntarily generate activity for remote locations in a natural (i.e. non-BMI) task? Here, we have provided evidence that remote replay is a strong candidate for deliberate, internal activation of remote plans.

## METHODS

### Experimental Details

#### Subjects

Adult male Long-Evans rats, weighing ∼300-400 g at the time of surgery, were individually housed in home cages fitted with custom-made running wheels throughout training and after surgery on a 12 h light / 12 h dark schedule. All procedures were performed according to the Janelia Research Campus Institutional Animal Care and Use Committee guidelines on animal welfare.

#### Chronic electrode implant surgery and targeting of hippocampal area CA1

Procedures were similar to (*51*, *58*). Animals were anesthetized with isoflurane and secured in a stereotaxic frame. The skull surface was cleaned and dried with hydrogen peroxide, then a custom 3D-printed ring was attached to the skull using dental adhesives (OptiBond Universal, Kerr) to form a base for the future implant enclosure. A small craniotomy was made above the cerebellum for a ground wire. Four elongated craniotomies were made over the dorsal hippocampus (centered approximately at AP –2.9 mm, ML ±2.2 mm and AP –4.2 mm, ML ±3.0 mm). The dura was softened with collagenase before probe insertion. Each of the four silicon probes (8-shank 64-site Buzsaki 64 or 6-shank 64-site BuzsakiSP 64 probe, Neuronexus) was attached to a screw-based Nano Drive (Ronal Tool) and successively lowered using a stereotaxic holder into the brain to a depth of 1.0-1.5 mm beneath the surface and cemented to the base ring. (If a probe did not go in smoothly, additional dura was manually opened.) Once all probes were in the brain, artificial dura was applied to cover the craniotomies. Each probe was connected to two 32-channel analog headstages (Intan Technologies), and the headstages remained with the animal inside the implant. All of the wiring was connected to a custom PCB so that there was a single connector that needed to be attached to for recording. A custom 3D-printed shell with silver conductive paint and strips of copper mesh covering the inside and attached to the ground (for shielding) was attached to the base ring. In addition, for purposes not related to this study, two animals (rats 1 and 2) had two more probes each implanted in the medial prefrontal cortex (mPFC) (AP +3.5 mm, ML ±0.5 mm) and, in one animal (rat 3), an opsin-containing viral construct was injected into the hippocampus in a previous procedure and optical fibers were attached to the silicon probes.

#### Maze apparatus

Figure 1A shows a to-scale diagram of the maze and its components. Briefly, there are three long parallel arms coming off of a central hub area. The hub area contains a touchpad on the floor that is sensitive to the weight of the animal in order to detect the rats’ presence in the area. Also located in the hub area is a running wheel (10-cm width, 29.5-cm in diameter, Lafayette Instrument, 8086W) attached to an optical motion detector and brake mechanism (HB6M-500-500-I-S-D, US Digital). The brake mechanism was used to constrain the wheel to turn in only one (and the same) direction for all trials. The entrance to each arm from the hub area could be either opened or closed using automatically controlled hydraulic doors. Each maze arm is 15.25 cm wide at the base and the walls are angled outward, reaching a vertical height of 33 cm. Each arm is equipped with an inner and outer beam break (Omron E3F2-R4C4-P1) as well as a water port at the end that has an optical lick detector. Each water reward port consisted of a custom 3D-printed housing for the lick detector attached to a Harvard Apparatus PHD-2000 syringe pump for reward delivery. The entire maze system was controlled using an Arduino Mega 2560-based board. Behavioral events such as timestamped records of beam breaks, touchpad, wheel speed, opening and closing of doors, reward delivery, reward port licks, and tally of correct and error trials was saved into an event file.

#### Behavioral training

Rats were trained using the following method after having silicon probes implanted into the brain. After being given sufficient time to recover from the implant surgery, the rat was started on acclimation to the maze and then training on the task. The two days of maze acclimation went as follows: on the first day, the rats were introduced to the maze by being placed inside the hub area with all doors open and were allowed to freely explore and run in the wheel. On the second day, the rat was placed in the maze but the doors were opened and closed with close supervision by the experimenter to acclimate the rat to the noise of the doors. After these two days, the rat was started on water restriction such that reward from the maze was the rats’ only source of water. The day after water restriction was started, the rat was placed in the hub area with all arm doors closed. The door to the left arm was opened following a 0.5-second wheel run above the speed threshold (18.5 cm/s). Once the door opened the rat was allowed to explore the left arm and the experimenter made sure the rat was aware that the reward port was dispensing water. Then the rat was allowed to return to the hub area where the door closed behind him. Rats quickly learned in 1-2 trials that turning the wheel caused a door to open. During this training period, only a single door opened each trial and the doors opened in what would be the correct order for the final task: left, center, right, center. The rat was given time in the maze every day to perform this “forced” version of the task that simply required running in the wheel to get a door to an arm to open, running down the arm to get reward, and returning to the hub area to start the next trial. Over time, the required amount of time for continuous running in the wheel above the speed threshold was gradually increased each day by 0.5 or 1 s (if the speed dropped below the threshold before the minimum duration, the timer was restarted). Typically, at the start of each day the rat ran the bare minimum amount in the wheel learned from the previous day (for example, 2 s) and found that this caused nothing to happen. The rat would then try again until eventually continuously running the amount of time to reach the duration needed for the day (for example, 2.5 s) to cause the door to open. The rat would then collect water reward at the end of the arm and go on to the next trial. Typically, the next trial would also involve multiple wheel run attempts at first, just short of the new time requirement; however, the rats quickly learned over a couple of trials the new required continuous wheel run duration and typically ran the required time for the majority of the day’s new trials. Once a rat was performing forced trials with a 6 s-long wheel run, it was moved on to the full task. This was simply done by starting the day with the maze set to the full task parameters: 8 s wheel run, followed by opening all three door arms after the run. A choice was registered by crossing a beam break near the end of an arm, at which point the two other arm doors closed to prevent access to the unchosen arms. If the selected arm was correct in the sequence, a water reward was delivered. Each reward consisted of 0.1 mL of water delivered slowly at a flow rate of 0.7 mL/min. Incorrect arm choices resulted in no water reward being delivered, thus over many days the rats slowly learned the required correct behavioral procedure sequence of left, center, right, center by trial and error. If an incorrect choice was made (e.g. incorrect left instead of correct right), in the next trial the animal needed to make the originally correct choice (e.g. right) in order to be correct and get reward–that is, an incorrect choice did not reset the correct arm sequence. Each rat had one behavioral session per day. Animals typically performed ∼90 trials over a total of ∼60-75 minutes. Note that for each day’s recording procedure, the rat was taken out of the cage and fed peanut butter treats in the experimenters’ lap while the implant’s top cover was removed and the electrophysiology recording cable and tracking LEDs were attached to the implant. Then the rat was placed into the hub area with one more peanut butter treat to eat while the experimenter returned to the recording system and started the recording of all behavioral parameters and electrophysiology data. Sessions were typically ended when the rats’ behavior seemed to indicate lack of motivation to continue, with the exception of attempts to make sure the rat did enough trials in a single day to get sufficient water.

#### Behavioral Electrophysiology Recording

Neural recording and tracking of position were carried out using the Amplipex recording system (Amplipex KJE-1001, www.amplipex.com). The Amplipex system is integrated with Intan multiplexing headstages (www.intantech.com/index.html) which permit recording from large numbers of channels over a thin light cable to minimize interference with behavior in freely moving rats. The commutator for recording was either from Doric Lenses (ERJ 12 HDMI-B2) or custom-made from Dragonfly (www.dragonflyinc.com). Neural activity was recorded at 20 kHz. Behavioral location within the maze was recorded by tracking LEDs on the rat’s implant at 30 Hz using a webcam integrated with the Amplipex system. A sync pulse from the maze behavioral control system was recorded on the Amplipex system in order to guarantee accurate syncing of behavioral information and neural activity recorded by Amplipex.

## Data Analysis

Analysis was performed using MATLAB. To compare a sample proportion to a chance level, we used the one-sample z-test for proportions (2-tailed). To compare two independent sample proportions to each other, we used the two-sample z-test for proportions (2-tailed).

### Spike Sorting

Spike sorting was performed using the NDManager, KlustaKwik, and Klusters suites. Spikes were detected from high pass filtered data by keeping events above a threshold of the mean +1.5 SD. Then the spike waveforms were stored and automatically sorted using the KlustaKwik suite. After automatic sorting, clustering was further refined manually using Klusters (*59*). All sorted units were included in the analysis (including all putative pyramidal cells and interneurons, with no minimum threshold for spatial information).

### Place and episode field maps

Place field maps were computed using the following methods. All place field maps were computed using periods when the rat was moving over 5 cm/s. Two-dimensional place fields were calculated using 2 cm x 2 cm bins. The binned spike count was divided by binned occupancy in order to obtain the binned firing rate, then the firing rate was smoothed with a Gaussian kernel with standard deviation (SD) of 2.75 cm. As place cells in the maze showed strong directional selectivity, therefore for each trial the outbound and inbound segments were labeled and treated separately (here outbound refers to the trajectory preceding reward collection, consisting of the rat moving away from the central hub area and out towards the end of the maze arm, and inbound for the return direction). For instance, inbound place fields were computed using activity during the return period from the reward port at the end of a given maze arm back to the hub area. For calculating linearized place fields, the X-Y tracking coordinates were projected onto the closest point on a path drawn down the middle of each arm, then the linear location was determined as the index along the line. Linearized place fields were calculated using the linearized coordinates and they were calculated separately for the inbound and outbound segments of each trial (the length of each arm was divided into 52 bins and the firing rate was smoothed using a Gaussian kernel with SD = 1 bin). Thus, each neuron had a 2D place field map as well as linearized place field maps for the inbound and outbound trajectories of each of the three arms, for a total of six linearized maps. Episode field maps for the wheel running period were computed with the following methods. The episode field map (one for all wheel runs over each session) was computed by analyzing the spike times relative to the beginning of each wheel run. Activity was binned into 125 ms bins, and the firing rate was smoothed using a Gaussian kernel with SD = 1 bin).

### Population Burst Event detection

To identify candidate replay events, we first detected population burst events (PBEs) (short periods of coordinated, high-frequency neural activity) during periods when the animal was located at the end of the maze arms. The summed spiking activity from all recorded neurons was binned into 1 ms time bins and smoothed with a Gaussian kernel with SD = 20 ms, yielding the population firing rate (PFR). For each trial, a threshold was calculated using the mean and SD of the PFR during the period at the arm end and when the rat’s speed was <5 cm/s. Successive times when the PFR was greater than the mean + 2 SD were considered a candidate PBE, then the start and end of the event were refined to be the times when the PFR crossed below a lower threshold = (mean + 2 SD)/2. Events separated by <5 ms were merged, and only events lasting ≥65 ms were retained as PBEs. Finally, PBEs were retained for decoding if they occurred when the animal was at a maze arm end and the animal’s speed was below a threshold of 27.5 cm/s (this high value was used because the speed was computed at 30 Hz and was not smoothed, thus small head movements during licking could occasionally cause relatively high speed values, and PBEs are known to occur during reward consumption).

### Bayesian decoding

Memoryless Bayesian decoding of the rat’s location from neural activity was performed as previously described (*20*, *24*). Activity was decoded in sliding 20 ms windows computed every 5 ms. We performed Bayesian decoding separately for the linear and the 2D spatial maps. For the linear map decoding, decoding was performed simultaneously across all six linear maps plus the wheel map, such that the probability across all seven maps summed to 1 in each 20 ms window.

### Replay detection

We searched for replay in each PBE. For each PBE, we took all the 20 ms time windows (at 5 ms steps) that occurred during that PBE. For the linear maps, for each window we only kept the decoded location (or time for wheel map) with maximum probability across all six arm maps and the wheel map (e.g. if the maximum occurred for the left arm outbound map, the corresponding location in that map was kept, and no other locations in the other maps were kept for that time window). Furthermore, the decoded location for a window was kept only if the probability for that location was >0.01, and the total number of active neurons in that window was 4 or more. This yielded seven maps in which to look for potential replay among the qualified decoded locations. In addition, because some replays have been found to be a mixture of different directional maps (*20*, *26*), we separately considered three non-directional maps (one for each arm) from the 2D spatial map. For this, we kept the maximum probability location for each window using the full 2D map (again, we kept it only if the probability was >0.01 and at least 4 neurons were active), then that location was projected (i.e. linearized) onto the nearest of three lines representing the left, center, and right arms (binned into one of 100 bins per arm). These three maps were added to the seven previous maps, then we performed the following steps to look for replay in any of the maps. For each map, we computed the weighted Spearman correlation (corval) of location versus rank-ordered time (modified from (*26*)) using all of the qualified decoded locations in that map. We then applied a threshold for the required minimum number of qualified decoded locations (numstep) in each map. This was done by first excluding from the count any decoded locations that were stationary across adjacent time windows (i.e. in the same spatial bin as in the previous window). That is, if a qualified decoded location in window n+1 was at the same location as a qualified decoded location in window n, then window n+1 was excluded from the count. Next, a map was considered as having a putative replay if the absolute value of corval was ≥0.8 and numstep was ≥8 (these are the values that were used for all the results in the main text, but all results were found to be robust to a range of corval and numstep values as illustrated in the supplementary materials). If a PBE had putative replays for two maps associated with the same arm (e.g. left outbound and 2D left arm), then the event was classified as a replay event associated with the map that had the highest absolute corval. In rare cases, if a PBE had a putative replays for multiple maps corresponding to different arms, or for the wheel, then the event was discarded. Finally, we classified each replay event in the following ways: Left/Center/Right (based on the arm map the replay was associated with), Local/Remote (local if the current location is in the same arm as the replay, remote if in a different arm), Inbound/Outbound (depending on the direction of propagation of the replay), and, if the replay was associated with one of the six original linear maps, Forward/Reverse (forward if the replay was associated with an outbound arm map and propagated outbound, or was associated with an inbound arm map and propagated inbound, and reverse otherwise).

### Remote replay-associated behavior analysis

For assessing the extent to which remote replays at the center arm predicted the arm choice in the next trial, we considered all remote outbound replays (note that almost all remote replays were outbound) which occurred when the animal was at the center arm end and it was the correct choice, and for which the previous arm choice was also correct (in order to avoid any ambiguity for the animal about which arm should be the next correct choice). For assessing the extent to which remote replays at an edge arm predicted the next arm choice, we considered all remote outbound replays which occurred when the animal was at an edge arm end, and it was the correct choice (note that the previous choice did not need to be correct since from an edge arm the center arm is unambiguously the next correct choice).

### Behavioral performance

The measurement of task performance (% trials in which the arm was chosen correctly, Fig. 1C, left) included all trials. For the task performance for trials beginning at an edge versus the center arm end (Fig. 1C, right), only trials starting from an edge arm in which the current trial was correct (i.e. the animal received a reward at that edge arm end) were included, and only trials starting from the center arm in which the current and previous trial were correct were included. These conditions were applied so that the next correct arm was the next arm in the standard sequence, which is the center arm from an edge arm (which requires 1-back memory), and the correct edge arm from the center arm (which requires 2-back memory). Otherwise, because the correct arm after an error is the arm the animal should have originally chosen, this edge-to-center versus center-to-edge trial comparison would not have been a pure measure of performance under 1-back versus 2-back memory conditions.

## ACKNOWLEDGMENTS

We thank A. Karpova and J. Magee for valuable advice and discussions; T. Tabachnik and the Instrument Design & Fabrication facility at Janelia Research Campus, A. Lustig, and the members of the Janelia Vivarium team for technical assistance; and M. Nardin and L. Grima for valuable comments on the manuscript.

## AUTHOR CONTRIBUTIONS

Conceptualization: B.L., A.K.L., S.R. Task design: B.L. Experimental apparatus and processing pipeline: E.P., Y.W. Experiments: B.L. Data analysis: B.L., A.K.L., S.R. Writing: B.L., A.K.L. Supervision: A.K.L., E.P., S.R.

## FUNDING

This work was supported by the Howard Hughes Medical Institute.

**Figure S1.**
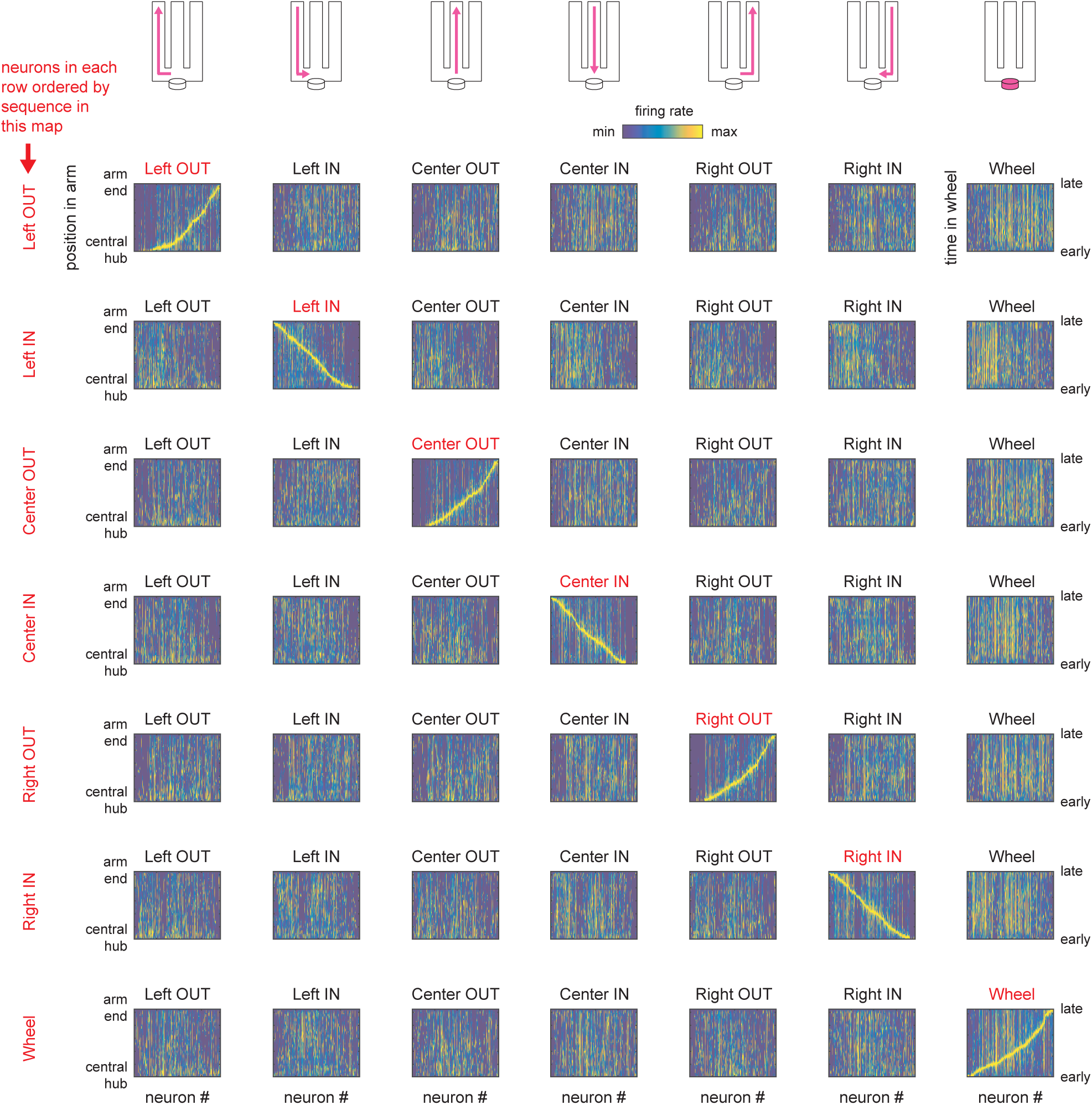
Firing rate maps from an example session. The seven linear maps are shown in each row, where for each row the neurons are ordered by the location of their peak firing rate in a given map (denoted in red). (If a neuron was silent in a given map, it was placed as having a peak at the central hub end of the arm.) The color scale for the firing rate is applied to each neuron separately.

**Figure S2.**
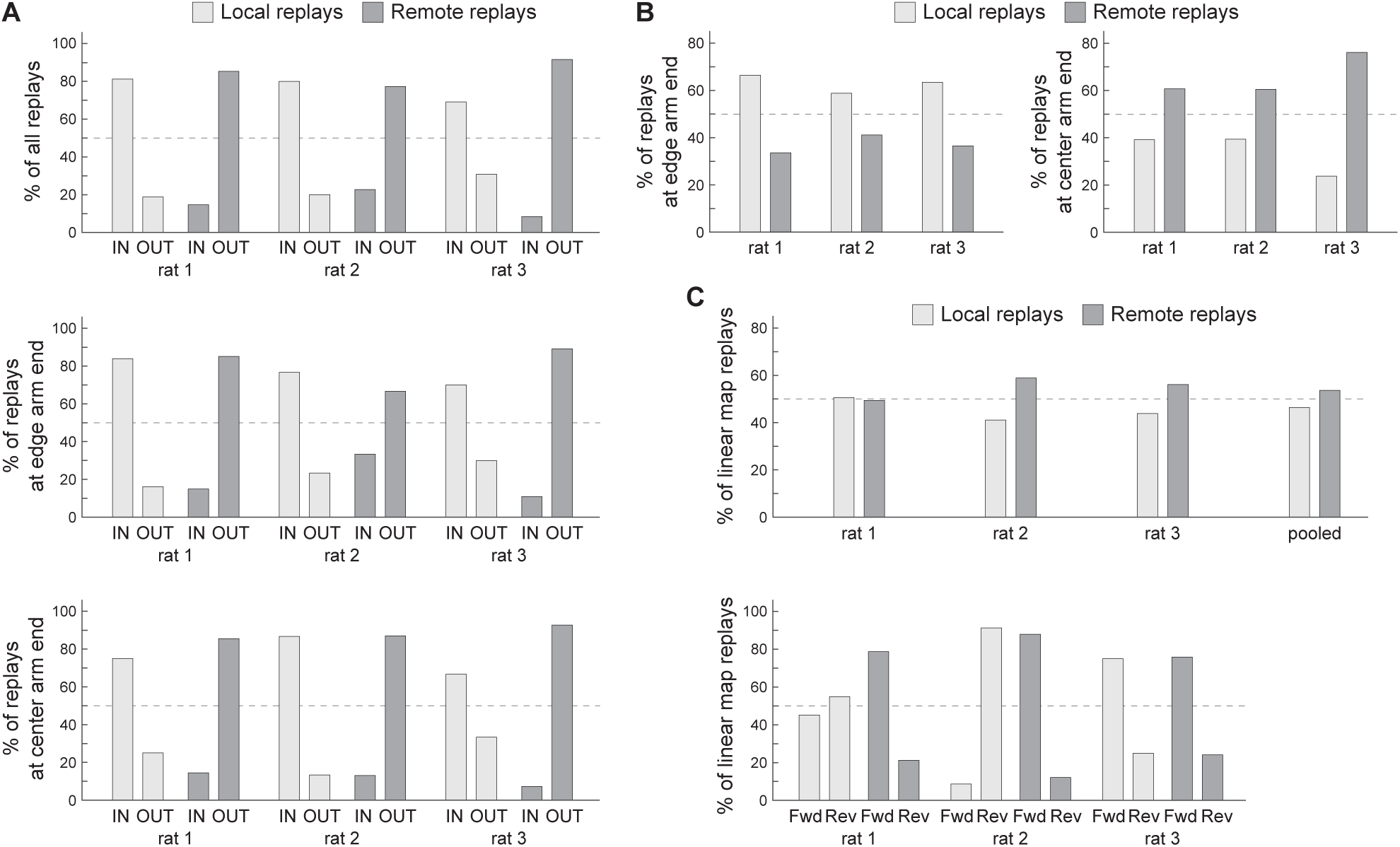
Individual animal replay proportions (related to Figure 2G-I). (A) The trends for local replays being strongly biased toward inbound trajectories and remote replays being strongly biased toward outbound trajectories is present in individual animals for all replays (top) as well as for replays while the animal is at an edge arm end (middle) or the center arm end (bottom). (A)-(C) The trends for each animal match the pooled results in Fig. 2G-I, except that the non-significant difference between forward versus reverse direction local replays in Fig. 2H is represented by trends in opposing directions in individual animals (C, bottom).

**Figure S3.**
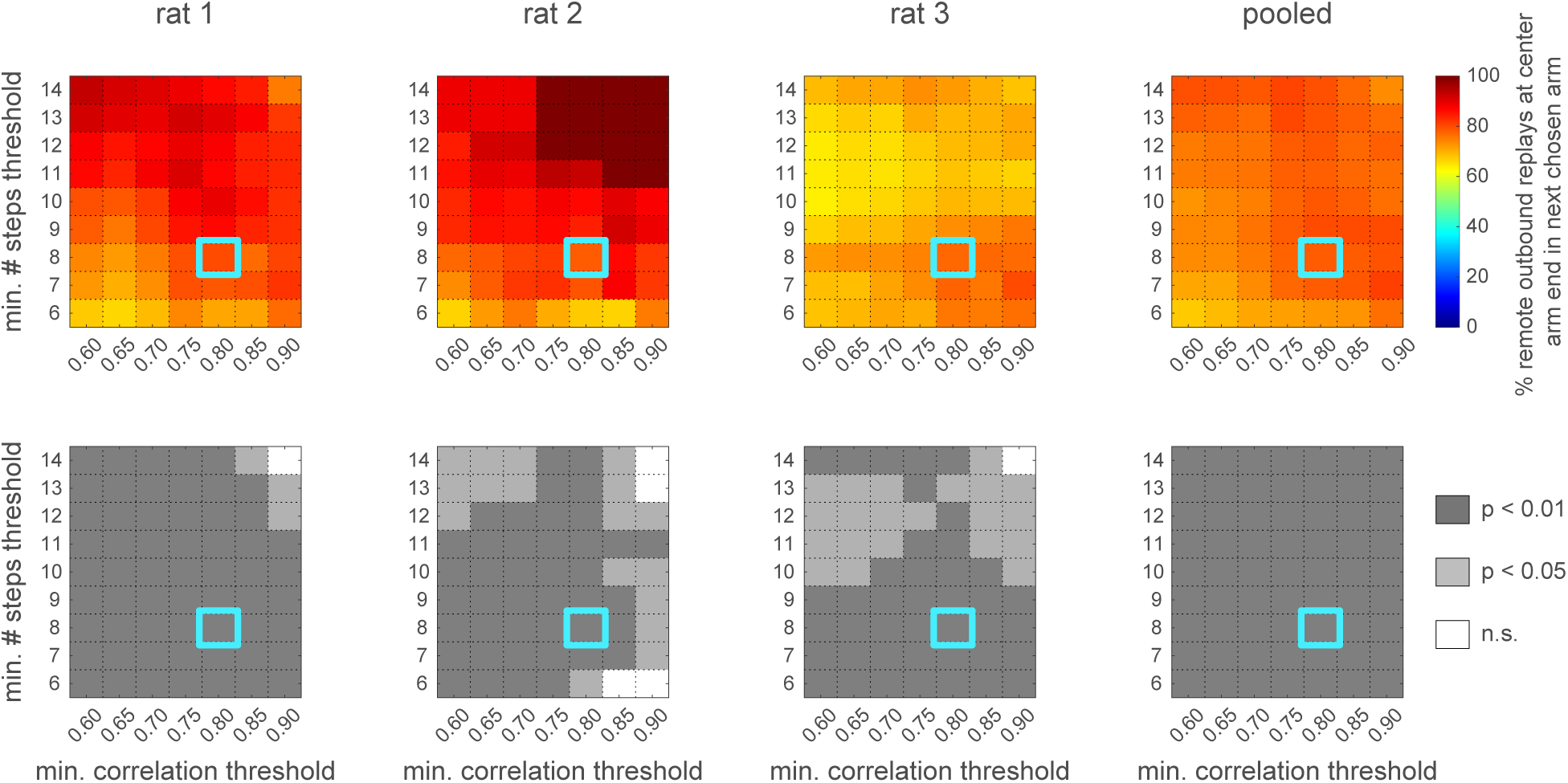
Remote outbound replays at the center arm end are strongly biased toward the animal’s next arm choice (related to Figure 3C). The bias and its significance are robust across a wide range of parameter values used for detecting replay events. Parameters used for the results in the main text and figures are outlined in cyan (numsteps = 8; corval = 0.8).

**Movie S1.** Example left-to-center arm trial from the three-arm delayed sequence (TADS) task. 1x speed.

**Movie S2.** Example remote replay while rat is at the center arm end that predicts the next chosen arm. 1x speed except for period when remote replay occurs (time at top). During the replay event, location is decoded in 20 ms windows stepped every 5 ms, with the posterior probability of the decoder shown for each window, then overlaid at the end. Note that an 8-second wheel run occurs between this replay and the next arm choice.

## REFERENCES

1. E. Tulving, W. Donaldson, Eds., Organization of Memory (Academic Press, San Diego, CA, 1972).

2. R. S. Rosenbaum, S. Köhler, D. L. Schacter, M. Moscovitch, R. Westmacott, S. E. Black, F. Gao, E. Tulving, The case of K.C.: contributions of a memory-impaired person to memory theory. Neuropsychologia 43, 989–1021 (2005).

3. W. B. Scoville, B. Milner, Loss of recent memory after bilateral hippocampal lesions. J. Neurol. Neurosurg. Psychiatry 20, 11–21 (1957).

4. D. L. Schacter, D. R. Addis, R. L. Buckner, Remembering the past to imagine the future: the prospective brain. Nat. Rev. Neurosci. 8, 657–661 (2007).

5. T. Suddendorf, M. C. Corballis, Mental time travel and the evolution of the human mind. Genet. Soc. Gen. Psychol. Monogr. 123, 133–167 (1997).

6. E. Tulving, “Episodic memory and autonoesis: Uniquely human?” in The Missing Link in CognitionOrigins of Self-Reflective Consciousness (Oxford University Press, 2005), pp. 3–56.

7. D. Hassabis, D. Kumaran, E. A. Maguire, Using imagination to understand the neural basis of episodic memory. J. Neurosci. 27, 14365–14374 (2007).

8. E. C. Tolman, Cognitive maps in rats and men. Psychol. Rev. 55, 189–208 (1948).

9. J. O’Keefe, L. Nadel, The Hippocampus as a Cognitive Map (Oxford University Press, London, England, 1978).

10. J. O’Keefe, J. Dostrovsky, The hippocampus as a spatial map. Preliminary evidence from unit activity in the freely-moving rat. Brain Res. 34, 171–175 (1971).

11. E. Pastalkova, V. Itskov, A. Amarasingham, G. Buzsáki, Internally generated cell assembly sequences in the rat hippocampus. Science 321, 1322–1327 (2008).

12. C. J. MacDonald, K. Q. Lepage, U. T. Eden, H. Eichenbaum, Hippocampal “time cells” bridge the gap in memory for discontiguous events. Neuron 71, 737–749 (2011).

13. W. E. Skaggs, B. L. McNaughton, Replay of neuronal firing sequences in rat hippocampus during sleep following spatial experience. Science 271, 1870–1873 (1996).

14. A. K. Lee, M. A. Wilson, Memory of sequential experience in the hippocampus during slow wave sleep. Neuron 36, 1183–1194 (2002).

15. G. Girardeau, K. Benchenane, S. I. Wiener, G. Buzsáki, M. B. Zugaro, Selective suppression of hippocampal ripples impairs spatial memory. Nat. Neurosci. 12, 1222–1223 (2009).

16. V. Ego-Stengel, M. A. Wilson, Disruption of ripple-associated hippocampal activity during rest impairs spatial learning in the rat. Hippocampus 20, 1–10 (2010).

17. I. Gridchyn, P. Schoenenberger, J. O’Neill, J. Csicsvari, Assembly-specific disruption of hippocampal replay leads to selective memory deficit. Neuron 106, 291–300.e6 (2020).

18. D. J. Foster, M. A. Wilson, Reverse replay of behavioural sequences in hippocampal place cells during the awake state. Nature 440, 680–683 (2006).

19. K. Diba, G. Buzsáki, Forward and reverse hippocampal place-cell sequences during ripples. Nat. Neurosci. 10, 1241–1242 (2007).

20. T. J. Davidson, F. Kloosterman, M. A. Wilson, Hippocampal replay of extended experience. Neuron 63, 497–507 (2009).

21. M. P. Karlsson, L. M. Frank, Awake replay of remote experiences in the hippocampus. Nat. Neurosci. 12, 913–918 (2009).

22. A. C. Singer, L. M. Frank, Rewarded outcomes enhance reactivation of experience in the hippocampus. Neuron 64, 910–921 (2009).

23. A. S. Gupta, M. A. A. van der Meer, D. S. Touretzky, A. D. Redish, Hippocampal replay is not a simple function of experience. Neuron 65, 695–705 (2010)

24. B. E. Pfeiffer, D. J. Foster, Hippocampal place-cell sequences depict future paths to remembered goals. Nature 497, 74–79 (2013).

25. A. C. Singer, M. F. Carr, M. P. Karlsson, L. M. Frank, Hippocampal SWR activity predicts correct decisions during the initial learning of an alternation task. Neuron 77, 1163–1173 (2013).

26. X. Wu, D. J. Foster, Hippocampal replay captures the unique topological structure of a novel environment. J. Neurosci. 34, 6459–6469 (2014).

27. H. F. Ólafsdóttir, C. Barry, A. B. Saleem, D. Hassabis, H. J. Spiers, Hippocampal place cells construct reward related sequences through unexplored space. Elife 4 (2015).

28. R. E. Ambrose, B. E. Pfeiffer, D. J. Foster, Reverse replay of hippocampal place cells is uniquely modulated by changing reward. Neuron 91, 1124–1136 (2016).

29. H. F. Ólafsdóttir, F. Carpenter, C. Barry, Task demands predict a dynamic switch in the content of awake hippocampal replay. Neuron 96, 925–935.e6 (2017).

30. C.-T. Wu, D. Haggerty, C. Kemere, D. Ji, Hippocampal awake replay in fear memory retrieval. Nat. Neurosci. 20, 571–580 (2017).

31. A. A. Carey, Y. Tanaka, M. A. A. van der Meer, Reward revaluation biases hippocampal replay content away from the preferred outcome. Nat. Neurosci. 22, 1450–1459 (2019).

32. H. Xu, P. Baracskay, J. O’Neill, J. Csicsvari, Assembly responses of hippocampal CA1 place cells predict learned behavior in goal-directed spatial tasks on the radial eight-arm maze. Neuron 101, 119–132.e4 (2019).

33. B. Bhattarai, J. W. Lee, M. W. Jung, Distinct effects of reward and navigation history on hippocampal forward and reverse replays. Proc. Natl. Acad. Sci. U. S. A. 117, 689–697 (2020).

34. A. K. Gillespie, D. A. Astudillo Maya, E. L. Denovellis, D. F. Liu, D. B. Kastner, M. E. Coulter, D. K. Roumis, U. T. Eden, L. M. Frank, Hippocampal replay reflects specific past experiences rather than a plan for subsequent choice. Neuron 109, 3149–3163.e6 (2021).

35. H. Igata, Y. Ikegaya, T. Sasaki, Prioritized experience replays on a hippocampal predictive map for learning. Proc. Natl. Acad. Sci. U. S. A. 118, e2011266118 (2021).

36. C. Zheng, E. Hwaun, C. A. Loza, L. L. Colgin, Hippocampal place cell sequences differ during correct and error trials in a spatial memory task. Nat. Commun. 12, 3373 (2021).

37. A. Berners-Lee, T. Feng, D. Silva, X. Wu, E. R. Ambrose, B. E. Pfeiffer, D. J. Foster, Hippocampal replays appear after a single experience and incorporate greater detail with more experience. Neuron 110, 1829–1842.e5 (2022).

38. X. Mou, A. Pokhrel, P. Suresh, D. Ji, Observational learning promotes hippocampal remote awake replay toward future reward locations. Neuron 110, 891–902.e7 (2022).

39. J. Widloski, D. J. Foster, Flexible rerouting of hippocampal replay sequences around changing barriers in the absence of global place field remapping. Neuron 110, 1547–1558.e8 (2022).

40. S. P. Jadhav, C. Kemere, P. W. German, L. M. Frank, Awake hippocampal sharp-wave ripples support spatial memory. Science 336, 1454–1458 (2012).

41. M. S. Nokia, J. E. Mikkonen, M. Penttonen, J. Wikgren, Disrupting neural activity related to awake-state sharp wave-ripple complexes prevents hippocampal learning. Front. Behav. Neurosci. 6, 84 (2012).

42. A. Fernández-Ruiz, A. Oliva, E. Fermino de Oliveira, F. Rocha-Almeida, D. Tingley, G. Buzsáki, Long-duration hippocampal sharp wave ripples improve memory. Science 364, 1082–1086 (2019).

43. W.-C. Jiang, S. Xu, J. T. Dudman, Hippocampal representations of foraging trajectories depend upon spatial context. Nat. Neurosci. 25, 1693–1705 (2022).

44. L. Deceuninck, F. Kloosterman, Disruption of awake sharp-wave ripples does not affect memorization of locations in repeated-acquisition spatial memory tasks. Elife 13 (2024).

45. W. Yang, C. Sun, R. Huszár, T. Hainmueller, K. Kiselev, G. Buzsáki, Selection of experience for memory by hippocampal sharp wave ripples. Science 383, 1478–1483 (2024).

46. G. Buzsáki, Hippocampal sharp wave-ripple: A cognitive biomarker for episodic memory and planning. Hippocampus 25, 1073–1188 (2015).

47. D. J. Foster, Replay comes of age. Annu. Rev. Neurosci. 40, 581–602 (2017).

48. H. R. Joo, L. M. Frank, The hippocampal sharp wave-ripple in memory retrieval for immediate use and consolidation. Nat. Rev. Neurosci. 19, 744–757 (2018).

49. H. F. Ólafsdóttir, D. Bush, C. Barry, The role of hippocampal replay in memory and planning. Curr. Biol. 28, R37–R50 (2018).

50. M. A. A. van der Meer, D. Bendor, Awake replay: off the clock but on the job. Trends Neurosci. 48, 257–267 (2025).

51. Y. Wang, S. Romani, B. Lustig, A. Leonardo, E. Pastalkova, Theta sequences are essential for internally generated hippocampal firing fields. Nat. Neurosci. 18, 282–288 (2015).

52. M. A. Wilson, B. L. McNaughton, Reactivation of hippocampal ensemble memories during sleep. Science 265, 676–679 (1994).

53. Z. Nádasdy, H. Hirase, A. Czurkó, J. Csicsvari, G. Buzsáki, Replay and time compression of recurring spike sequences in the hippocampus. J. Neurosci. 19, 9497–9507 (1999).

54. C. S. Mallory, J. Widloski, D. J. Foster, The time course and organization of hippocampal replay. Science 387, 541–548 (2025).

55. S. Romani, M. Tsodyks, Short-term plasticity based network model of place cells dynamics. Hippocampus 25, 94–105 (2015).

56. C. Lai, S. Tanaka, T. D. Harris, A. K. Lee, Volitional activation of remote place representations with a hippocampal brain-machine interface. Science 382, 566–573 (2023).

57. M. E. Coulter, A. K. Gillespie, J. Chu, E. L. Denovellis, T. T. K. Nguyen, D. F. Liu, K. Wadhwani, B. Sharma, K. Wang, X. Deng, U. T. Eden, C. Kemere, L. M. Frank, Closed-loop modulation of remote hippocampal representations with neurofeedback. Neuron 113, 949–961.e3 (2025).

58. M. Vandecasteele, S. Royer, M. Belluscio, A. Berényi, K. Diba, S. Fujisawa, A. Grosmark, D. Mao, K. Mizuseki, J. Patel, E. Stark, D. Sullivan, B. Watson, G. Buzsáki, Large-scale recording of neurons by movable silicon probes in behaving rodents. J. Vis. Exp., doi: 10.3791/3568-v (2012).

59. L. Hazan, M. Zugaro, G. Buzsáki, Klusters, NeuroScope, NDManager: a free software suite for neurophysiological data processing and visualization. J. Neurosci. Methods 155, 207–216 (2006).

